# Structural variability, expression profile and pharmacogenetics properties of TMPRSS2 gene as a potential target for COVID-19 therapy

**DOI:** 10.1101/2020.06.20.156224

**Authors:** Aleksei Zarubin, Vadim Stepanov, Anton Markov, Nikita Kolesnikov, Andrey Marusin, Irina Khitrinskaya, Maria Swarovskaya, Sergey Litvinov, Natalia Ekomasova, Murat Dzhaubermezov, Nadezhda Maksimova, Aitalina Sukhomyasova, Olga Shtygasheva, Elza Khusnutdinova, Magomed Radjabov, Vladimir Kharkov

## Abstract

The human serine protease TMPRSS2 gene is involved in the priming of the novel severe acute respiratory syndrome coronavirus 2 (SARS-CoV-2) proteins being one of the possible targets for COVID-19 therapy. TMPRSS2 gene is possibly co-expressed with SARS-CoV-2 cell receptor genes ACE2 and BSG, but only TMPRSS2 demonstrates tissue-specific expression in alveolar cells according to single cell RNA sequencing data. Our analysis of the structural variability of the TMPRSS2 gene based on genome-wide data of 76 human populations demonstrates that functionally significant missense mutation in exon 6/7 in TMPRSS2 gene, was found in many human populations in relatively high frequency, featuring region-specific distribution patterns. The frequency of the missense mutation encoded by the rs12329760, which previously was found to be associated with prostate cancer, is ranged between 10% and 63% being significantly higher in populations of Asian origin compared to European populations. In addition to SNPs, two copy numbers variants (CNV) were detected in the TMPRSS2 gene. Number of microRNAs have been predicted to regulate TMPRSS2 and BSG expression levels, but none of them is enriched in lung or respiratory tract cells. Several well studied drugs can downregulate the expression of TMPRSS2 in human cells, including Acetaminophen (Paracetamol) and Curcumin. Thus TMPRSS2 interaction with the SARS-CoV-2, its structural variability, gene-gene interactions, and expression regulation profiles, and pharmacogenomics properties characterize this gene as a potential target for COVID-19 therapy.

## Introduction

Novel severe acute respiratory syndrome coronavirus 2 (SARS-CoV-2) caused a pandemic of coronavirus diseases (COVID-19) and lead to a global public health crisis. Infection of the human cells with viral particles occurs through the binding of S viral proteins to the receptors of the host cell and their subsequent priming with proteases. ACE2 is considered the classic receptor for SARS-CoV-2, but there is evidence that it can also use the BSG receptor (CD147) [33]. The priming of viral proteins is carried out by the TMPRSS2 protease. No specific therapy has yet been developed for SARS-CoV-2. But it was found that blockers of all three proteins can prevent cell infection [15].

In addition to protein blocking, there are other mechanisms that can alter the expression level or affinity of the interaction of viral particles with specific proteins. Possible ways of expression differentiation include alteration of protein structure due to genetic variants (SNV, INDEL), copy number variation (CNV), variants affecting regulatory regions (eQTL), and epigenetic regulation (methylation, miRNA).

TMPRSS2 gene in humans encodes a transmembrane protein that belongs to the serine protease family. Alternatively spliced transcript variants encoding different isoforms have been found for this gene. TMPRSS2 protein involved in prostate carcinogenesis via overexpression of ETS transcription factors, such as ERG and ETV1, through gene fusion. TMPRSS2-ERG gene fusion, present in 40% - 80% of prostate cancers in humans, is one of the molecular subtypes that has been associated with predominantly poor prognosis [25, 28].

TMPRSS2 protease proteolytically cleaves and activates glycoproteins of many viruses including spike proteins of human coronavirus 229E (HCoV-229E) and human coronavirus EMC (HCoV-EMC); the fusion glycoproteins of Sendai virus (SeV), human metapneumovirus (HMPV), human parainfluenza 1, 2, 3, 4a and 4b viruses (HPIV) [1, 4, 13–15, 29, 30]. It has been shown that both the SARS coronavirus of 2003 severe acute respiratory syndrome outbreak in Asia (SARS-CoV) and the SARS-CoV-2 are activated by TMPRSS2 and can thus be inhibited by TMPRSS2 inhibitors [15]. In this work, we report the data on genetic variability of TMPRSS2 gene in 76 human populations of North Eurasia in comparison with worldwide populations; analyze the data on expression and its regulation of TMPRSS2 gene, its interaction with SARS-CoV-2 receptors, and its pharmacogenetic properties.

## Materials and Methods

### Structural variability data

Allele frequency for worldwide populations were downloaded from GnomAD database containing information on the frequencies of genomic variants from more than 120 thousand exomes and 15 thousand of whole genomes [18]. These data were used to search SNVs and INDELs in TMPRSS2 gene. Data on copy number variation were obtained from CNV Control Database [21].

Data on allele frequencies in 76 populations of North Eurasia were extracted from the unpublished own dataset of population genomics data obtained by genotyping using Illumina Infinium genome-wide microarrays. In brief, 1836 samples from 76 human populations were genotyped for 1748250 SNVs and INDELs using Infinium Multi-Ethnic Global-8 Kit. Populations represent various geographic regions of North Eurasia (Eastern Europe, Caucasus, Central Asia, Siberia, North-East Asia) and belong to various linguistic families (Indo-European, Altaic, Uralic, North Caucasian, Chukotko-Kamchatkan, Sino-Tibetan., Yeniseian). DNA samples were collected under informed consent and deposited to DNA bank of the Research Institute for Medical Genetics, Tomsk National Medical Research Center, Tomsk, Russia and DNA bank of the Institute of Biochemistry and Genetics, Ufa Federal Research Centre of the Russian Academy of Sciences. The study was approved by the Ethical Committee of the Research Institute for Medical Genetics, Tomsk National Medical Research Center. Data on 4 missense mutations in TMPRSS2 gene were extracted from the dataset. CNV search was performed using Markov model algorithm for high-resolution copy number variation detection in whole-genome SNP implemented in PennCNV tool [34].

To determine possible functional impact of detected SNVs, the Polymorphism Phenotyping v2 (Poly-Phen-2) tool was used [2]. Poly-Phen estimates the impact of the mutation on the stability and function of the protein using the structural and evolutionary analyses of the amino acid substitution. The tool evaluates the probability of the mutation to be probably damaging, possibly damaging, benign or of unknown significance using quantitative prediction with a score.

### Bioinformatics analysis of gene expression, miRNA intercaction and pharmacogenomics

Analysis of protein – protein interactions of SARS-CoV-2 interacting proteins was carried out using the GeneMANIA and STRING web resources [32, 38]. Single cell RNA sequencing data were downloaded from the PanglaoDB database which contains more than 1300 single cell sequencing samples [12]. Lung cells single cell RNA-seq data were obtained from the Sequence Read Archive (SRA) [22] and processed in R software environment using the Seurat package [31].

Analysis of the interaction of miRNAs with target proteins was performed using information from two databases, miRTarBase, which contains information from more than 8000 referenced sources about experimentally confirmed micro RNA - protein interactions [16], and miRPathDB database containing experimentally confirmed and predicted miRNA-protein interactions [19]. Data on the differential expression of miRNAs in various cell cultures were downloaded from the database of the FANTOM5 project [10]. DRUGBANK database [35] was used to search for the drugs which may change the level of protein expression.

## Results and Discussion

### Protein-protein interaction network of SARS-CoV-2 interacting genes

Protein-protein interaction networks obtained with two different tools (GeneMANIA and STRING) (Fig. 1) demonstrates that TMPRSS2 is co-expressing with other SARS-CoV-2 interacting genes, despite showed contradictory co-expression patterns. According to GeneMANIA, TMPRSS2 is co-expressed with BSG, while STRING indicates co-expression between ACE2 and TMPRSS2. Interestingly that BSG shows the maximum number of protein-protein interactions in both networks.

**Figure 1.**
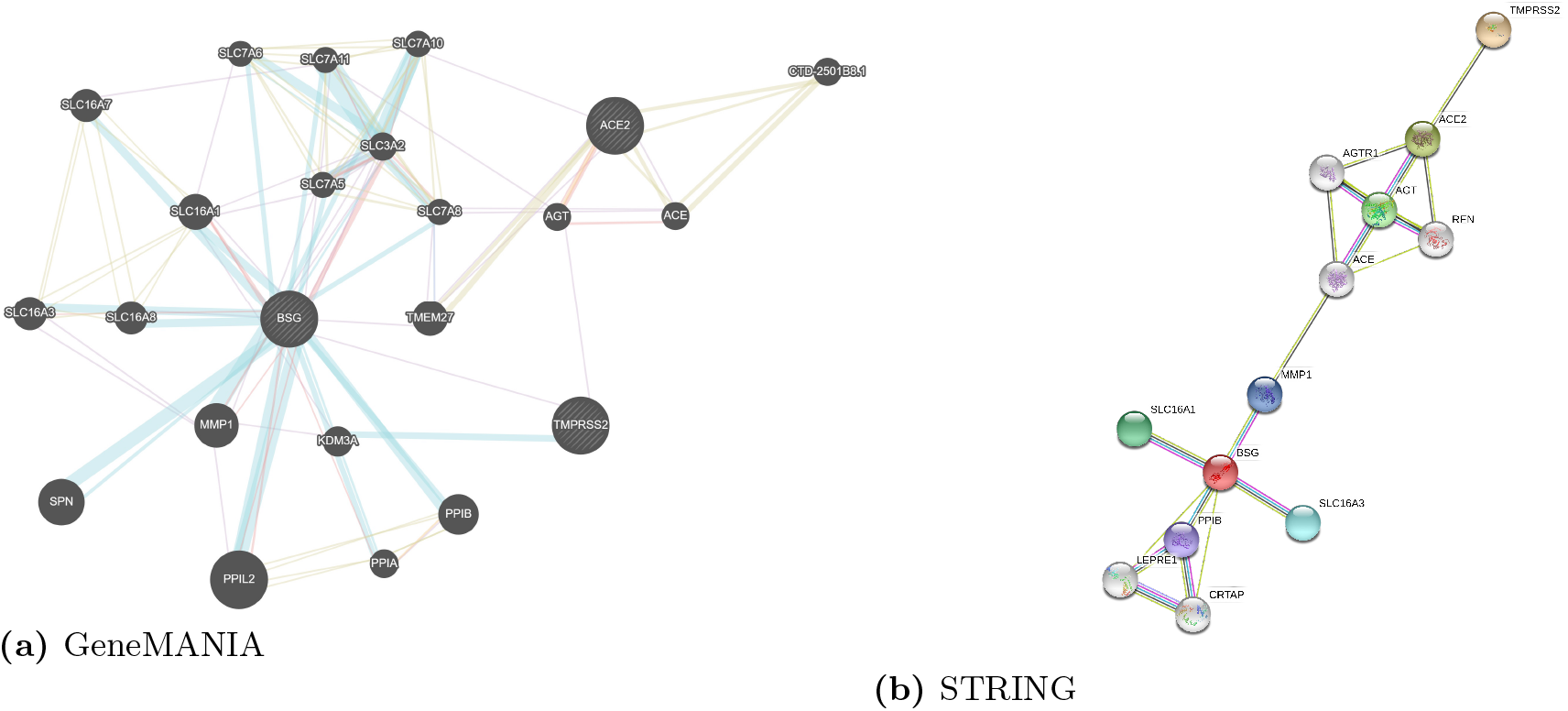
Protein-protein interaction network of SARS-CoV-2 interacting genes

Basigin, or extracellular matrix metalloproteinase inducer (EMMPRIN), also known as cluster of differentiation 147 (CD147), encoded by the BSG gene, is a transmembrane glycoprotein that belongs to the immunoglobulin superfamily and is a determinant for the Ok blood group system. BSG protein plays an important role in targeting the monocarboxylate transporters SLC16A1, SLC16A3, SLC16A8, SLC16A11 and SLC16A12 to the plasma membrane [7, 24, 27]. BSG is involved in spermatogenesis, embryo implantation, neural network formation and tumor progression. It stimulates adjacent fibroblasts to produce matrix metalloproteinases (MMPS). BSG seems to be a receptor for oligomannosidic glycans and according to in vitro experiments can promote outgrowth of astrocytic processes [7, 24, 27]. BSG is which is involved in tumor development, plasmodium invasion and virus infection [8, 17, 23, 26, 36, 37]. Previous data on severe acute respiratory syndrome indicate that BSG plays a functional role in facilitating SARS-CoV invasion for host cells, and CD147-antagonistic peptide-9 has a high binding rate to HEK293 cells and an inhibitory effect on SARS-CoV [9]. Based on the similarity of SARS-CoV and SARS-CoV-2, the function of BSG in invasion for host cells by SARS-CoV-2 can be assumed. The exact role of BSG in COVID-19 is still unknown, but recently it was found that CD147 may bind spike protein of SARS-CoV-2 [33]. The preliminary data on a small sample of COVID-19 patients demonstrated that Meplazumab, a humanized anti-CD147 antibody, efficiently improved the recovery of patients with SARS-CoV-2 pneumonia with a favorable safety profile [6].

### Expression of ACE2, BSG and TMPRSS2 in single cells

Data on expression profiles of SARS-CoV-2-interacting genes in various tissues demonstrates that ACE2 has a high level of expression only in testicles (Fig. 2a). Highest expression of BSG was found in germ cells, endothelium of various localization, fibroblasts and some other cell types (Fig. 2b). TMPRSS2 showed a high level of expression in the prostate, intestines, and lungs (Fig. 2c).

**Figure 2.**
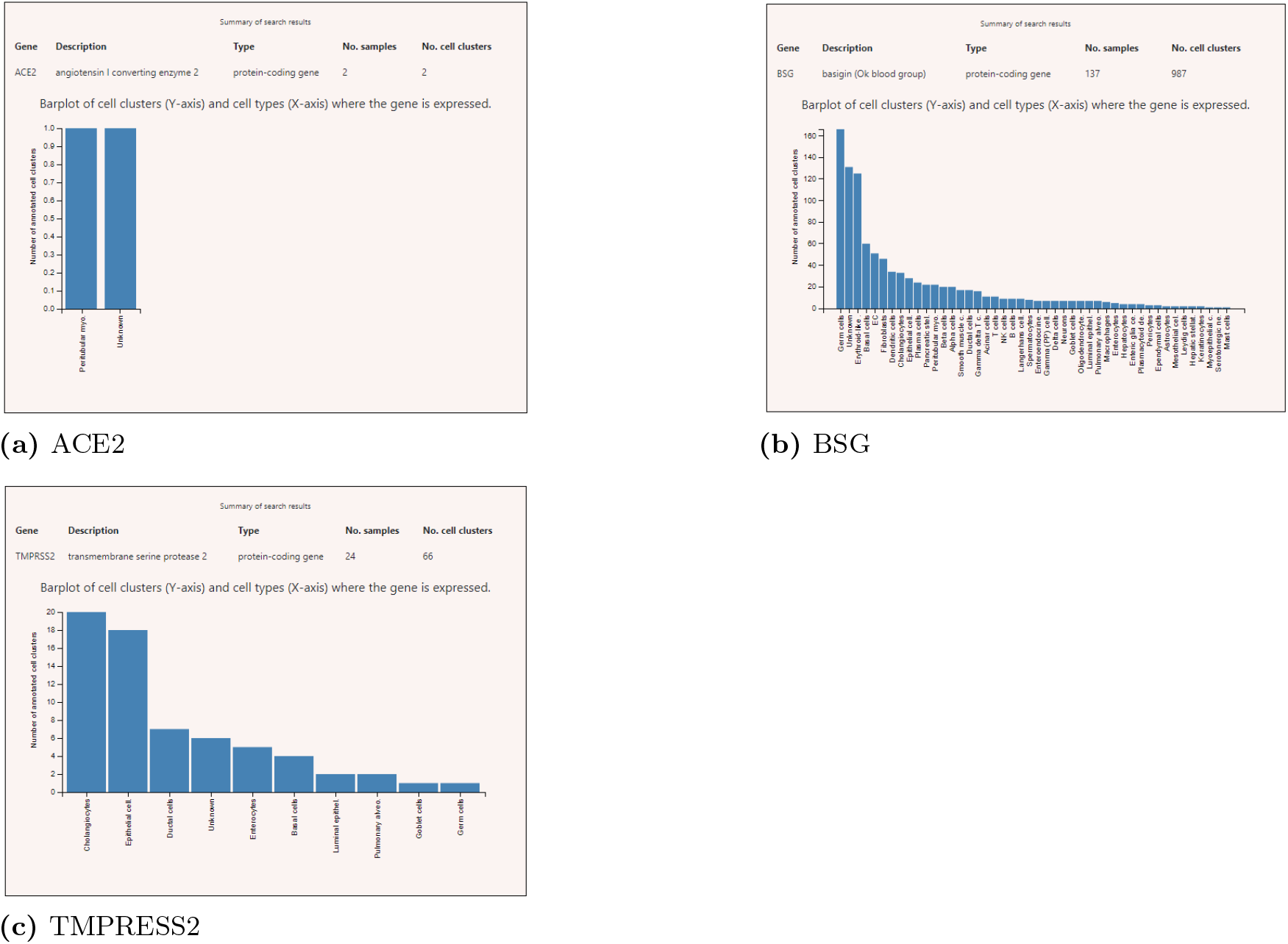
Expression level in various cells

In addition, the expression of these proteins was analyzed in a single sample (SRS2769051) of proximal stromal lung cells (Fig. 3). ACE2 had a low level of expression in pulmonary alveolar cells and as well as in fibroblasts. BSG is characterized by the average level of expression in fibroblasts and alveolar cells. Only TMPRSS2 gene demonstrates tissue-specific expression in alveolar cells.

**Figure 3.**
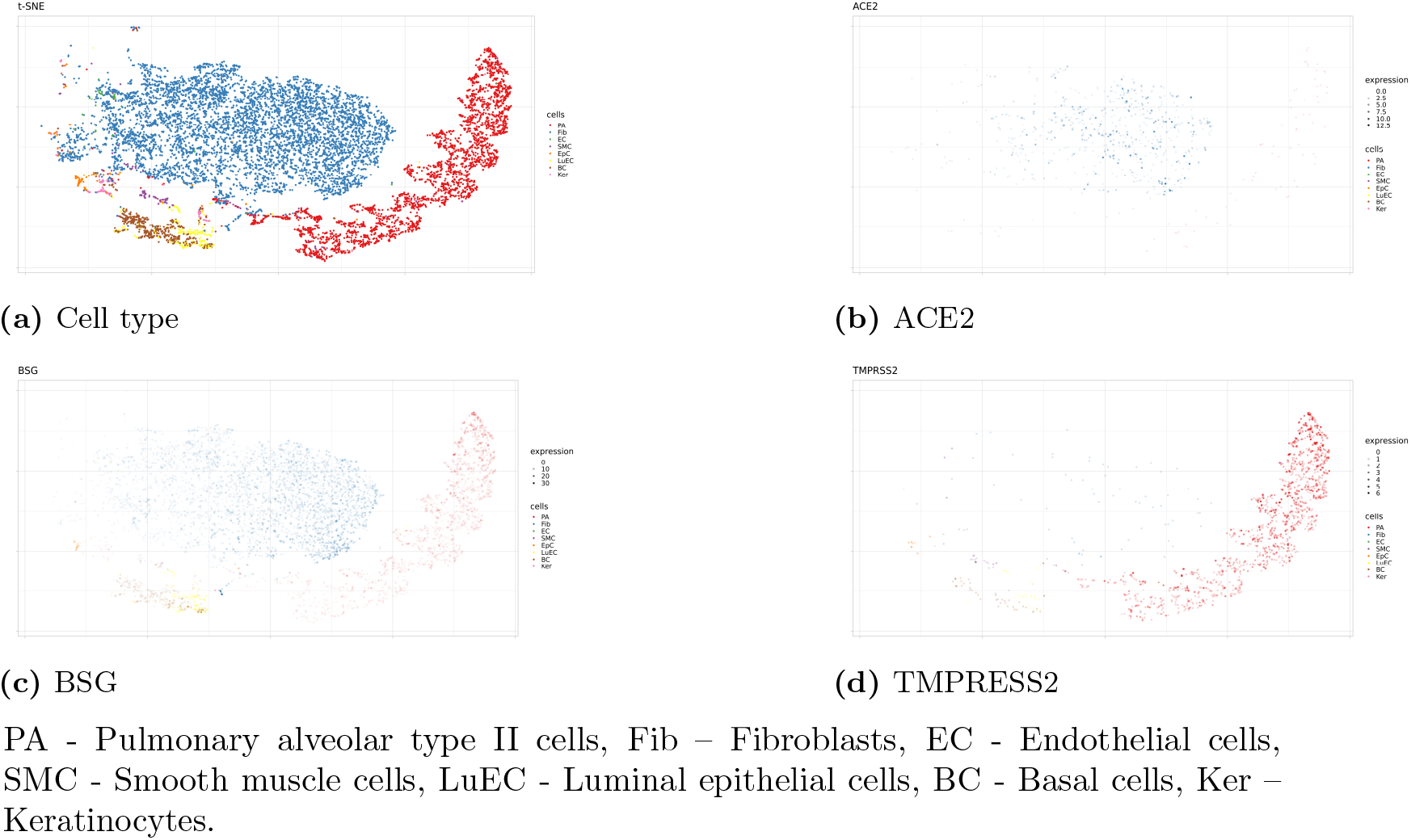
Expression level ACE2, BSG and TMPRESS2 in a single sample (SRS2769051) of proximal stromal lung cells.

Given the high specificity of expression of TMPRSS2 in lung, we further studied genomic and epigenomic properties of the gene that may possibly affect the level of its expression and the affinity of interaction with viral particles.

### SNV and INDEL variants or the TMPRSS2 gene

According information accumulated in GnomAD database, 1025 SNVs and INDELs of various frequencies, functional impact and localization have been described in TMPRSS2 gene. This list includes 332 missense variants, 17 frameshits, 64 splice sites variants, 14 stop codon mutations and 3 inframe INDELs. But among frequent variants (MAF > 0.01) there are only 13 intronic polymorphisms, 5 synonymous variants and 2 missense mutations (rs12329760 and rs75603675). Both missense variants have high frequencies (24.8% and 35.0% in gnomAD, respectively) (Table 1). The variant rs12329760 is the mutation of C to T in the position 589 of the gene that led to change of valine to methionine in the amino acid position 197 (exon 7) of transmembrane protease serine 2 isoform 1, or in position 160 (exon 6) of isoform 2. This mutation is predicted by Poly-Phen-2 to be probably damaging with a score of 0.997 (sensitivity: 0.41; specificity: 0.98). The allele T of the TMPRSS2 rs12329760, was positively associated with TMPRSS2-ERG fusion by translocation and also was associated with increased risk for prostate cancer in European and Indian populations [5, 11]. The rs75603675 (C to A transition in position 23, Gly8Val) was not reported to be associated with prostate cancer or with other clinical condition.

An interesting feature of both frequent missense variants is the difference in prevalence between European and Asian populations. Rs12329760 is 15% more frequent in populations of East Asia (38%) than in European populations (23%). For rs75603675 the difference is even more significant: minor allele reaches 42% in European populations and about 1% in populations of East Asia.

As to CNV, controlDB database contains only one deletion in TMPRSS2 gene (1 copy variant) with relatively low frequency (1.2%) (Table 2)

### Frequency of protein changing allelic variants of the TMPRSS2 gene in populations of North Eurasia

In order to study the population differentiation in TMPRSS2 functional variants in more details, we searched for TMPRSS2 allelic frequency in our own unpublished data on 76 populations of North Eurasia based on 1836 samples genotyped using genome-wide microarrays. Four missense mutations and two CNV in TMPRSS2 gene were found in our dataset. We summarized the frequency of TMPRSS2 missense mutations in North Eurasian population (Table 3) in comparison to worldwide data (Table 4). Three missense mutations (rs148125094, rs143597099, and rs201093031) were very rare variants, while rs12329760, previously associated with prostate cancer, were found with high frequency in all populations. The data on the second high-frequency missense variant in TMPRSS2 gene according to GnomAD database (s75603675) were not available because of the absence of this SNP in the microarray used in our study. The minor allele of variant rs148125094 was found on only 2 chromosomes (total frequency 0,00054) – in single heterozygous individuals from Karelian and Abkhaz populations. The variant rs143597099 was present only in one heterozygote from the Veps population. The variant rs201093031 was found in North-East Asian Nivkh and Udege populations with the frequency of 7%, and in a single Tuvan individual from Siberia. The frequency of the probably damaging minor allele of the rs12329760 polymorphism ranged between 10% (in Khvarshi population from Dagestan) and 63% (in Sagays Khakas). In general, the minor allele T has higher frequency in Siberia and Central Asia (both around 35%), while lowest frequency of the damaging variant was found in North Caucasus (19%), Dagestan (22%), and Eastern Europe (29%). This distribution correlates with the worldwide data demonstrating much higher frequency of the minor allele in Asian populations (36 – 41%) comparing to Europeans (22-24%) (see Table 4).

In addition, we detected CNV in 2 samples. In the first case, an increase in the number of copies covering the entire gene was found in the Karanogai individual. Second CNV was a affecting exons 3-7 fiund in a single Kumyk individual.

Thus potentially functionally significant variant in TMPRSS2 gene were found in many human populations in relatively high frequency, demonstrating region-specific distribution patterns. Both variants – probably damaging SNV and heterozygous deletion of the gene – may significantly contribute to the interaction of the human serine protease with the viral spike proteins changing the efficacy of the priming of viral proteins by the TMPRSS2 protease. However, the role of TMPRSS2 gene and its variants in the interaction with SARS-CoV-2 and in viral infectivity still needs to be elucidated.

### Regulation of expression of TMPRSS2

#### eQTLs

According to the GTEx Analysis V8 database, the TMPRSS2 gene contains 136 eQTLs (including 60 down-regulating and 76 up-regulating variants) that significantly alters its expression in lung tissues (Table 5). But in general, all these eQTLs have only minor effect on gene expression. The average slope of the regression line (value that characterized the strength of eQTL effect) is around 0.09 both for down- and up-regulating variants. The strongest single variant can change the expression by 13%.

#### miRNAs

According to the miRTarBase and miRPathDB databases, no experimentally proven miRNAs regulating TMPRSS2 were detected. It is worth to note that TMPRSS2 and BSG genes have the same predicted regulatory miRNAs.

Top 30 microRNAs predicted to regulate TMPRSS2 and BSG were analyzed for enrichment in various cell types using FANTOM5 database. None of the top miRNAs is enriched in lung or respiratory tract cells, but 3 miRNAs showed slight expression in immune and endothelial cells (Table. 6).

#### Pharmaco-transcriptomics of TMPRSS2

According to the DRUGBANK database, 9 drugs can reduce the level of expression of TMPRSS2. For 5 of them (Acetaminophen / Paracetamol, Curcumin, Cyclosporine, and Ethinylestradiol) this effect is clinically approved (Table 7). Information on the direction of the effect of Estradiol is conflicting – in different experiments it shows downregulation or upregulation effect on TMPRSS2 expression.

Two drugs from the list above (Acetaminophen / Paracetamol and Curcumin) were also considered as a possible therapy for COVID-19 [3]. Acetaminophen is currently being discussed as a possible drug for the correction of fever in patients with COVID-19. The discovered feature of this drug to reduce the level of expression of TMPRSS2 may be an additional argument in favor of its use, compared with other NSAIDs. Curcumin, a widely used food supplement, has the predicted ability to block Main Protease (Mpro) of SARS-CoV-2 [20], and may be studied further in relation to COVID-19 therapy.

## Conclusions

TMPRSS2 protein plays a crucial role in the process of SARS-CoV-2 activation in the human cells. The gene encoding this protease demonstrates high level of genetic variability as well as many variants which may regulate its expression levels. Despite very few potential functionally significant variants in the gene are of relatively high frequency, population-specific patterns of TMPRSS2 variability may contribute in some extent to the different viral infectivity of SARS-CoV-2 in populations of various geographic origins.

TMPRSS2 is probably co-expressed with SARS-CoV-2 receptors (ACE2 and BSG), but only the TMPRSS2 protease demonstrates tissue –specific expression an alveolar cell, target cell type for SARS-CoV-2 virus. It is an indication that TMPRSS2 is potentially the most promising target for COVID-19 therapy, based on the specific expression in lung, its important role in the process of cell infection and communication with other proteins involved in the infection process. Several well studied drugs can downregulate the expression of TMPRSS2 in human cells, including Acetaminophen (Paracetamol) and Curcumin. Both deserve close attention as possible anti-COVID-19 drugs due to their approved effects on TMPRSS2 expression, as well as because of a long history of their use, known side effects, and wide availability.

## Acknowledgments

This work is partially supported by Russian Foundation for Basic Research (project # 18-29-13045)

## Notes

### Competing Interest Statement

The authors have declared no competing interest.

## References

1. M. Abe, M. Tahara, K. Sakai, H. Yamaguchi, K. Kanou, K. Shirato, M. Kawase, M. Noda, H. Kimura, S. Matsuyama, et al. Tmprss2 is an activating protease for respiratory parainfluenza viruses. Journal of virology, 87(21):11930–11935, 2013.

2. I. Adzhubei, D. M. Jordan, and S. R. Sunyaev. Predicting functional effect of human missense mutations using polyphen-2. Current protocols in human genetics, 76(1):7–20, 2013.

3. W. Alhazzani, M. H. Møller, Y. M. Arabi, M. Loeb, M. N. Gong, E. Fan, S. Oczkowski, M. M. Levy, L. Derde, A. Dzierba, et al. Surviving sepsis campaign: guidelines on the management of critically ill adults with coronavirus disease 2019 (covid-19). Intensive care medicine, pages 1–34, 2020.

4. S. Bertram, R. Dijkman, M. Habjan, A. Heurich, S. Gierer, I. Glowacka, K. Welsch, M. Winkler, H. Schneider, H. Hofmann-Winkler, et al. Tmprss2 activates the human coronavirus 229e for cathepsin-independent host cell entry and is expressed in viral target cells in the respiratory epithelium. Journal of virology, 87(11):6150–6160, 2013.

5. A. Bhanushali, P. Rao, V. Raman, P. Kokate, A. Ambekar, S. Mandva, S. Bhatia, and B. Das. Status of tmprss2–erg fusion in prostate cancer patients from india: correlation with clinico-pathological details and tmprss2 met160val polymorphism. Prostate International, 6(4):145–150, 2018.

6. H. Bian, Z.-H. Zheng, D. Wei, Z. Zhang, W.-Z. Kang, C.-Q. Hao, K. Dong, W. Kang, J.-L. Xia, J.-L. Miao, et al. Meplazumab treats covid-19 pneumonia: an open-labelled, concurrent controlled add-on clinical trial. MedRxiv, 2020.

7. J. J. Castorino, S. M. Gallagher-Colombo, A. V. Levin, P. G. FitzGerald, J. Polishook, B. Kloeckener-Gruissem, E. Ostertag, and N. J. Philp. Juvenile cataract-associated mutation of solute carrier slc16a12 impairs trafficking of the protein to the plasma membrane. Investigative ophthalmology & visual science, 52(9):6774–6784, 2011.

8. A. P. V. Castro, T. M. Carvalho, N. Moussatché, and C. R. Damaso. Redistribution of cyclophilin a to viral factories during vaccinia virus infection and its incorporation into mature particles. Journal of virology, 77(16):9052–9068, 2003.

9. Z. Chen, L. Mi, J. Xu, J. Yu, X. Wang, J. Jiang, J. Xing, P. Shang, A. Qian, Y. Li, et al. Function of hab18g/cd147 in invasion of host cells by severe acute respiratory syndrome coronavirus. The Journal of infectious diseases, 191(5):755–760, 2005.

10. D. De Rie, I. Abugessaisa, T. Alam, E. Arner, P. Arner, H. Ashoor, G. Åström, M. Babina, N. Bertin, A. M. Burroughs, et al. An integrated expression atlas of mirnas and their promoters in human and mouse. Nature biotechnology, 35(9):872, 2017.

11. L. M. FitzGerald, I. Agalliu, K. Johnson, M. A. Miller, E. M. Kwon, A. Hurtado-Coll, L. Fazli, A. B. Rajput, M. E. Gleave, M. E. Cox, et al. Association of tmprss2-erg gene fusion with clinical characteristics and outcomes: results from a population-based study of prostate cancer. BMC cancer, 8(1):230, 2008.

12. O. Franzén, L.-M. Gan, and J. L. Björkegren. Panglaodb: a web server for exploration of mouse and human single-cell rna sequencing data. Database, 2019, 2019.

13. I. Glowacka, S. Bertram, M. A. Müller, P. Allen, E. Soilleux, S. Pfefferle, I. Steffen, T. S. Tsegaye, Y. He, K. Gnirss, et al. Evidence that tmprss2 activates the severe acute respiratory syndrome coronavirus spike protein for membrane fusion and reduces viral control by the humoral immune response. Journal of virology, 85(9):4122–4134, 2011.

14. A. Heurich, H. Hofmann-Winkler, S. Gierer, T. Liepold, O. Jahn, and Pöhlmann. Tmprss2 and adam17 cleave ace2 differentially and only proteolysis by tmprss2 augments entry driven by the severe acute respiratory syndrome coronavirus spike protein. Journal of virology, 88(2):1293–1307, 2014.

15. M. Hoffmann, H. Kleine-Weber, S. Schroeder, N. Krüger, T. Herrler, S. Erichsen, S. Schiergens, G. Herrler, N.-H. Wu, A. Nitsche, et al. Sars-cov-2 cell entry depends on ace2 and tmprss2 and is blocked by a clinically proven protease inhibitor. Cell, 2020.

16. H.-Y. Huang, Y.-C.-D. Lin, J. Li, K.-Y. Huang, S. Shrestha, H.-C. Hong, Y. Tang, Y.-G. Chen, C.-N. Jin, Y. Yu, et al. mirtarbase 2020: updates to the experimentally validated microrna–target interaction database. Nucleic acids research, 48(D1):D148–D154, 2020.

17. Q. Huang, J. Li, J. Xing, W. Li, H. Li, X. Ke, J. Zhang, T. Ren, Y. Shang, H. Yang, et al. Cd147 promotes reprogramming of glucose metabolism and cell proliferation in hcc cells by inhibiting the p53-dependent signaling pathway. Journal of hepatology, 61(4):859–866, 2014.

18. K. J. Karczewski, L. C. Francioli, G. Tiao, B. B. Cummings, J. Alföldi, Q. Wang, R. L. Collins, K. M. Laricchia, A. Ganna, D. P. Birnbaum, et al. Variation across 141,456 human exomes and genomes reveals the spectrum of loss-of-function intolerance across human protein-coding genes. BioRxiv, page 531210, 2019.

19. T. Kehl, F. Kern, C. Backes, T. Fehlmann, D. Stöckel, E. Meese, H.-P. Lenhof, and A. Keller. mirpathdb 2.0: a novel release of the mirna pathway dictionary database. Nucleic acids research, 48(D1):D142–D147, 2020.

20. S. Khaerunnisa, H. Kurniawan, R. Awaluddin, S. Suhartati, and S. Soetjipto. Potential inhibitor of covid-19 main protease (mpro) from several medicinal plant compounds by molecular docking study. Prepr. doi10, 20944:1–14, 2020.

21. A. Koike, N. Nishida, D. Yamashita, and K. Tokunaga. Comparative analysis of copy number variation detection methods and database construction. BMC genetics, 12(1):29, 2011.

22. R. Leinonen, H. Sugawara, M. Shumway, and I. N. S. D. Collaboration. The sequence read archive. Nucleic acids research, 39(suppl 1):D19–D21, 2010.

23. M. Lu, J. Wu, Z.-W. Hao, Y.-K. Shang, J. Xu, G. Nan, X. Li, Z.-N. Chen, and H. Bian. Basolateral cd147 induces hepatocyte polarity loss by e-cadherin ubiquitination and degradation in hepatocellular carcinoma progress. Hepatology, 68(1):317–332, 2018.

24. C. Manoharan, C. Manoharan, M. C. Wilson, C. Manoharan, M. C. Wilson, R. B. Sessions, and A. P. Halestrap. The role of charged residues in the transmembrane helices of monocarboxylate transporter 1 and its ancillary protein basigin in determining plasma membrane expression and catalytic activity. Molecular membrane biology, 23(6):486–498, 2006.

25. A. Pettersson, R. E. Graff, S. R. Bauer, M. J. Pitt, R. T. Lis, E. C. Stack, N. E. Martin, L. Kunz, K. L. Penney, A. H. Ligon, et al. The tmprss2: Erg rearrangement, erg expression, and prostate cancer outcomes: a cohort study and meta-analysis. Cancer Epidemiology and Prevention Biomarkers, 21(9):1497–1509, 2012.

26. T. Pushkarsky, G. Zybarth, L. Dubrovsky, V. Yurchenko, H. Tang, H. Guo, B. Toole, B. Sherry, and M. Bukrinsky. Cd147 facilitates hiv-1 infection by interacting with virus-associated cyclophilin a. Proceedings of the National Academy of Sciences, 98(11):6360–6365, 2001.

27. V. Rusu, E. Hoch, J. M. Mercader, D. E. Tenen, M. Gymrek, C. R. Hartigan, M. DeRan, M. von Grotthuss, P. Fontanillas, A. Spooner, et al. Type 2 diabetes variants disrupt function of slc16a11 through two distinct mechanisms. Cell, 170(1):199–212, 2017.

28. O. R. Saramäki, A. E. Harjula, P. M. Martikainen, R. L. Vessella, T. L. Tammela, and T. Visakorpi. Tmprss2: Erg fusion identifies a subgroup of prostate cancers with a favorable prognosis. Clinical cancer research, 14(11):3395–3400, 2008.

29. K. Shirato, M. Kawase, and S. Matsuyama. Middle east respiratory syndrome coronavirus infection mediated by the transmembrane serine protease tmprss2. Journal of virology, 87(23):12552–12561, 2013.

30. A. Shulla, T. Heald-Sargent, G. Subramanya, J. Zhao, S. Perlman, and T. Gallagher. A transmembrane serine protease is linked to the severe acute respiratory syndrome coronavirus receptor and activates virus entry. Journal of virology, 85(2):873–882, 2011.

31. T. Stuart, A. Butler, P. Hoffman, C. Hafemeister, E. Papalexi, W. M. Mauck III, Y. Hao, M. Stoeckius, P. Smibert, and R. Satija. Comprehensive integration of single-cell data. Cell, 177(7):1888–1902, 2019.

32. D. Szklarczyk, A. L. Gable, D. Lyon, A. Junge, S. Wyder, J. Huerta-Cepas, M. Simonovic, N. T. Doncheva, J. H. Morris, P. Bork, et al. String v11: protein–protein association networks with increased coverage, supporting functional discovery in genome-wide experimental datasets. Nucleic acids research, 47(D1):D607–D613, 2019.

33. K. Wang, W. Chen, Y.-S. Zhou, J.-Q. Lian, Z. Zhang, P. Du, L. Gong, Y. Zhang, H.-Y. Cui, J.-J. Geng, et al. Sars-cov-2 invades host cells via a novel route: Cd147-spike protein. BioRxiv, 2020.

34. K. Wang, M. Li, D. Hadley, R. Liu, J. Glessner, S. F. Grant, H. Hakonarson, and M. Bucan. Penncnv: an integrated hidden markov model designed for high-resolution copy number variation detection in whole-genome snp genotyping data. Genome research, 17(11):1665–1674, 2007.

35. D. S. Wishart, Y. D. Feunang, A. C. Guo, E. J. Lo, A. Marcu, J. R. Grant, T. Sajed, D. Johnson, C. Li, Z. Sayeeda, et al. Drugbank 5.0: a major update to the drugbank database for 2018. Nucleic acids research, 46(D1):D1074–D1082, 2018.

36. M.-Y. Zhang, Y. Zhang, X.-D. Wu, K. Zhang, P. Lin, H.-J. Bian, M.-M. Qin, W. Huang, D. Wei, Z. Zhang, et al. Disrupting cd147-rap2 interaction abrogates erythrocyte invasion by plasmodium falciparum. Blood, The Journal of the American Society of Hematology, 131(10):1111–1121, 2018.

37. P. Zhao, W. Zhang, S.-J. Wang, X.-L. Yu, J. Tang, W. Huang, Y. Li, H.-Y. Cui, Y.-S. Guo, J. Tavernier, et al. Hab18g/cd147 promotes cell motility by regulating annexin ii-activated rhoa and rac1 signaling pathways in hepatocellular carcinoma cells. Hepatology, 54(6):2012–2024, 2011.

38. K. Zuberi, M. Franz, H. Rodriguez, J. Montojo, C. T. Lopes, G. D. Bader, and Q. Morris. Genemania prediction server 2013 update. Nucleic acids research, 41(W1):W115–W122, 2013.

